## Supplementary Information for "Crystal structure of the human NKR-P1 bound to its lymphocyte ligand LLT1 reveals receptor clustering in the immune synapse"

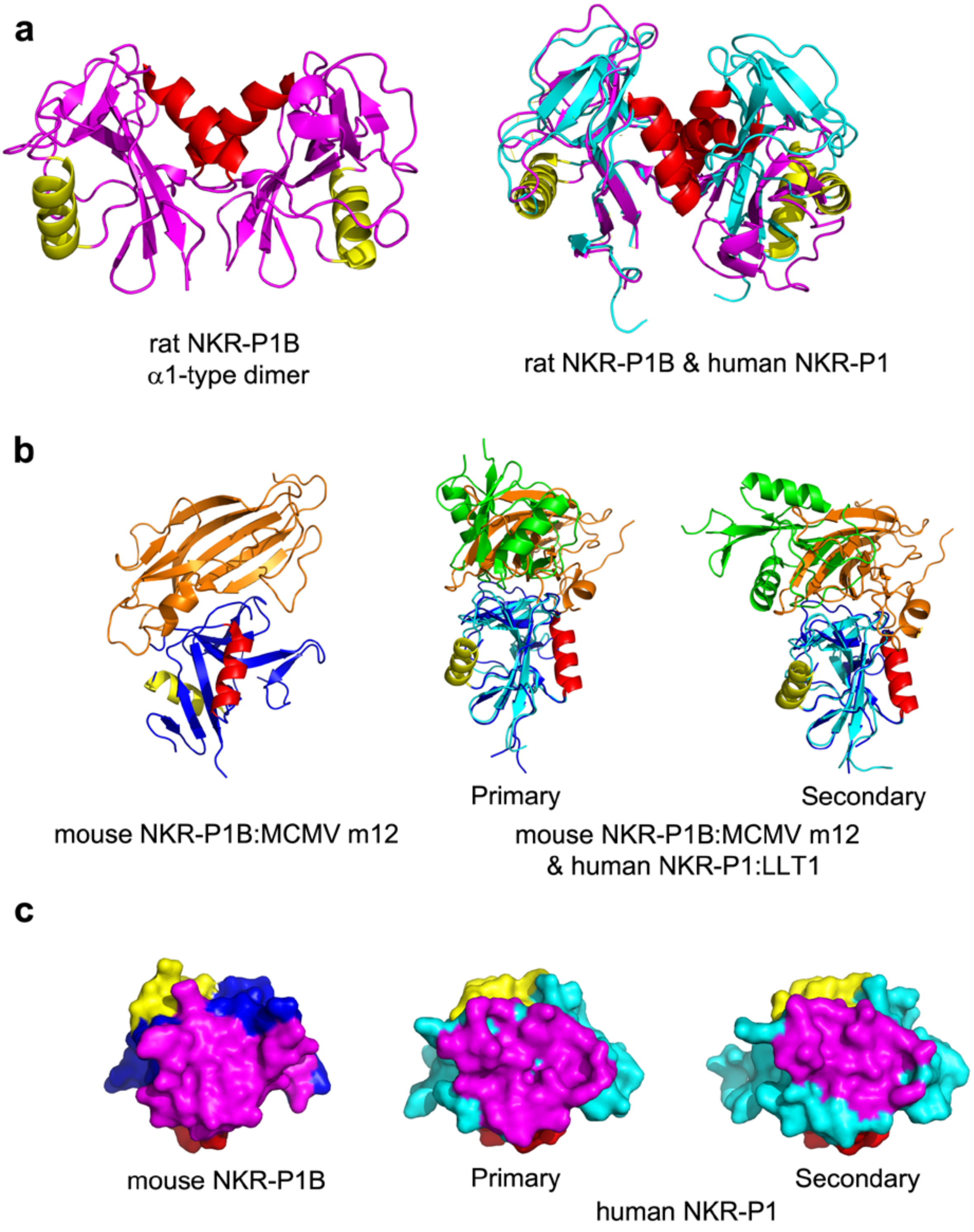

```
mouse NKR-P1B   90  AKLECPQDWLSHRDKCFHVSQVSNTWKECRIDCDKKGATLLLIQDQEELRFLLDSIKEKYNSFWIGLS
human NKR-P1    90  GLLNCPIYWQQLREKCLLFSHTVNPWNNSLADCSTKESSLLLIRDKDELIHTQNLIRDKAILFWIGLN

mouse NKR-P1B  158  YTLTDMNWKWINGTAFNSDVLKITGVTENGSCAAISGEKVTSEGCSSDNRWICQKELNHETPCNDS
human NKR-P1   158  FSLSEKNWKWINGSFLNSNDLEIRGDAKENSCISISQTSVYSEYCSTEIRWICQKELTPVRNKVYP
```

**Figure S1. Comparison of rat, murine and human NKR-P1 structures.** (**a**) Comparison between covalent disulfide dimer of rat NKR-P1B (PDB ID 5J2S, magenta) and non-covalent dimer of human NKR-P1 (PDB ID 5MGR, cyan); helices α1 and α2 are shown in red and yellow, respectively. Structural alignment of rat NKR-P1B and human NKR-P1 homodimers, prepared by aligning only one monomer from each dimer, is shown on the right-hand side. (**b**) Comparison between the structure of mouse NKR-P1B in complex with MCMV immunoevasin m12 (PDB ID 5TZN, blue and orange, respectively) and structure of human NKR-P1 in complex with LLT1 (PDB ID 5MGT, cyan and green, respectively) in primary and secondary interaction modes; helices α1 and α2 are shown in red and yellow, respectively. Structural alignment of the mouse NKR-P1B and human NKR-P1 monomers was used to compare their ligand-bound complexes, as shown on the right-hand side. (**c**) Interacting residues of NKR-P1 molecules shown in (b) are mapped on their respective surfaces in magenta while other colors follow the coding in (b); all molecules are oriented identically. The same residues are also highlighted in the sequence alignment below, in the yellow background for mouse NKR-P1B, and with the green background and purple triangles for the primary and secondary interaction mode of human NKR-P1, respectively.

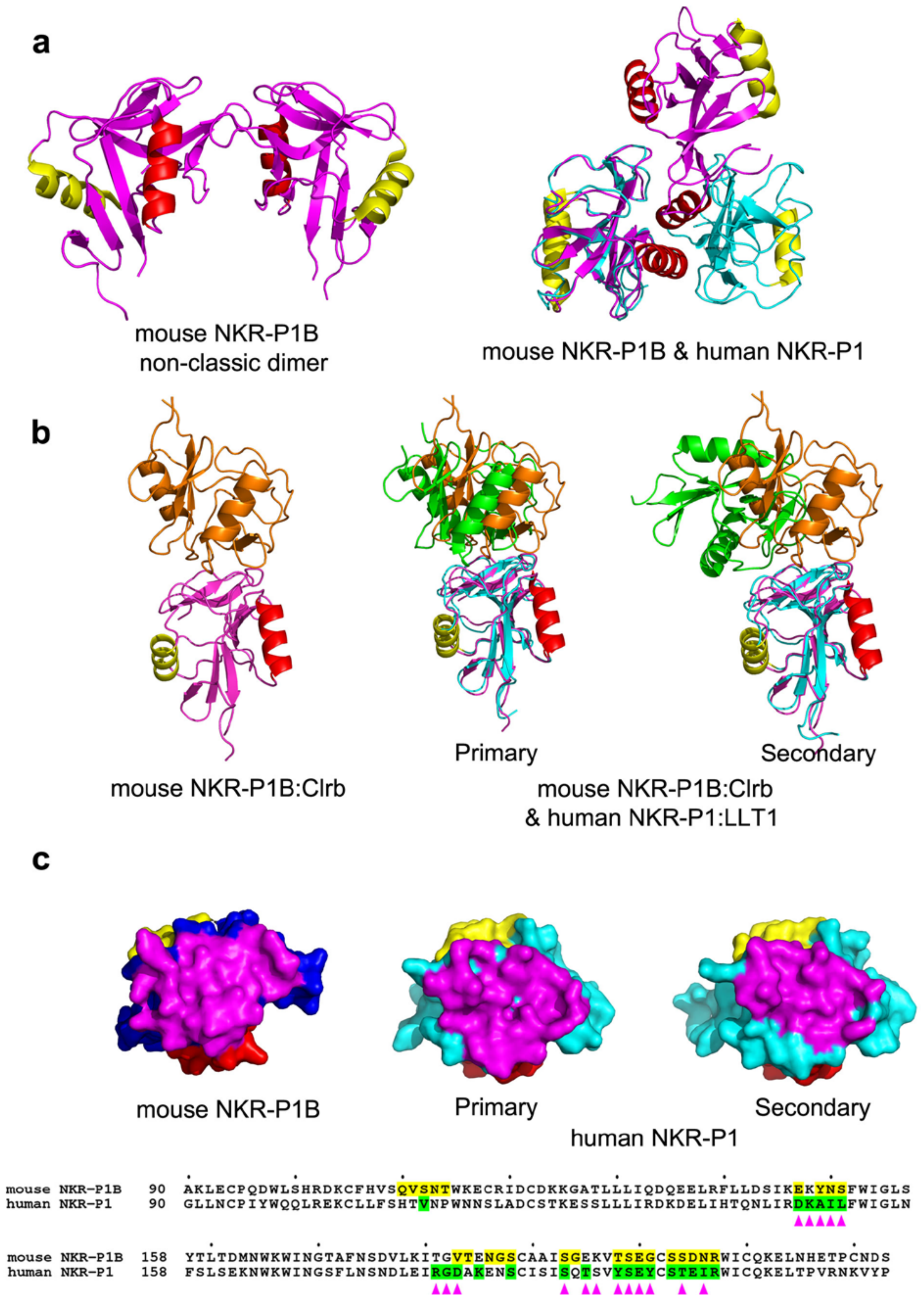

**Figure S2. Comparison of murine and human NKR-P1 complexes.** (**a**) Comparison between non-classical dimer of mouse NKR-P1B (PDB ID 6E7D, magenta) and non-covalent dimer of human NKR-P1 (PDB ID 5MGR, cyan); helices α1 and α2 are shown in red and yellow, respectively. Structural alignment of mouse NKR-P1B and human NKR-P1 homodimers, prepared by aligning only one monomer from each dimer, is shown on the right-hand side. (**b**) Comparison between the structure of mouse NKR-P1B in complex with Clrb (PDB ID 6E7D, magenta and orange, respectively) and structure of human NKR-P1 in complex with LLT1 (PDB ID 5MGT, cyan and green, respectively) in primary and secondary interaction modes; helices α1 and α2 are shown in red and yellow, respectively. Structural alignment of the mouse NKR-P1B and human NKR-P1 monomers was used to compare their ligand-bound complexes, as shown on the right-hand side. (**c**) Interacting residues of NKR-P1 molecules shown in (b) are mapped on their respective surfaces in magenta while other colors follow the coding in (b); all molecules are oriented identically. The same residues are also highlighted in the sequence alignment below, in the yellow background for mouse NKR-P1B, and with the green background and purple triangles for the primary and secondary interaction mode of human NKR-P1, respectively.

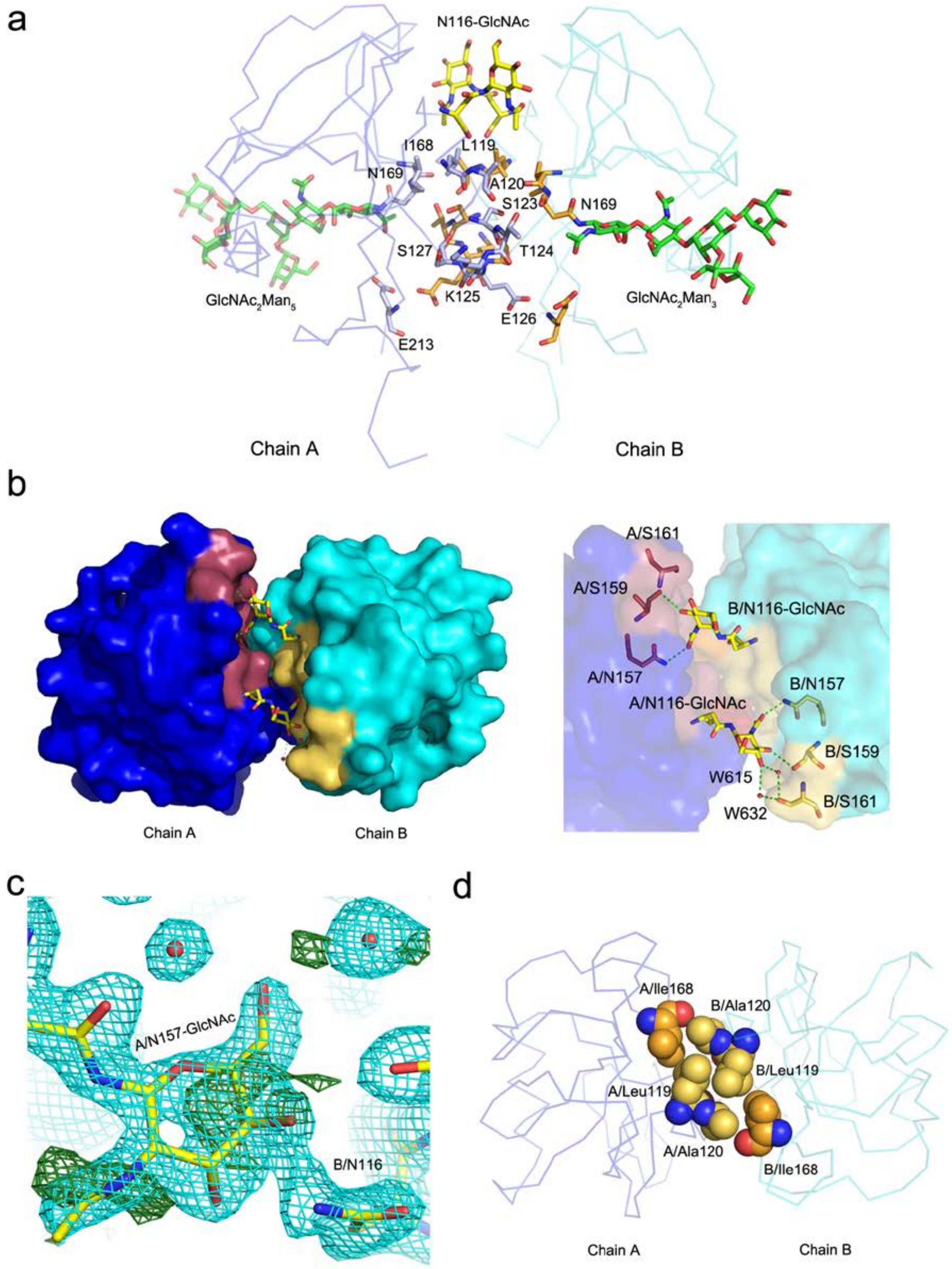

**Figure S3. Glycosylation affects the dimerization interface of human NKR-P1.** (**a**) Dimerization interface of human NKR-P1. Subunits of human NKR-P1 are shown as Cα-trace (blue and cyan), and the dimer contact residues are shown as sticks with carbon atoms colored in light blue (blue subunit) and orange (cyan subunit); for clarity, only the residues of the blue subunit are labeled. The first GlcNAc unit N-linked to Asn116 and the carbohydrate chain N-linked to Asn169, observable in the NKR-P1_glyco structure, are shown with carbon atoms colored yellow and green, respectively. (**b**) Top view of the dimerization interface. The NKR-P1 subunits surfaces are colored blue and cyan. The GlcNAc units bound to Asn116 are shown as sticks with carbon atoms in yellow. Contact residues between the GlcNAc bound to chain A, and the chain B, are shown in yellow, whereas contact residues between the GlcNAc bound to chain B, and the chain A, are shown in purple. Hydrogen bonds are shown as green-dashed lines with a detailed view on the right-hand side. (**c**) Mixed glycosylation states at the dimer interface in the NKR-P1_deglyco structure. The GlcNAc unit N-linked to Asn157 of chain A is modeled with occupancy of 0.5, while the second GlcNAc unit present at Asn116 of chain B is not modeled. Contours of $2mF_o$-$DF_c$ (2.8 σ, cyan) and $mF_o$-$DF_c$ (1 σ, green) electron density maps are shown. (**d**) Small hydrophobic core in the central part of the NKR-P1 dimerization interface (subunits colored as in (a)). The central residues are shown as spheres with carbon atoms in yellow. The carbon atoms of Ille168 residues (whose mutation decreases the ability of NKR-P1 to bind LLT1)[1] are shown in orange.

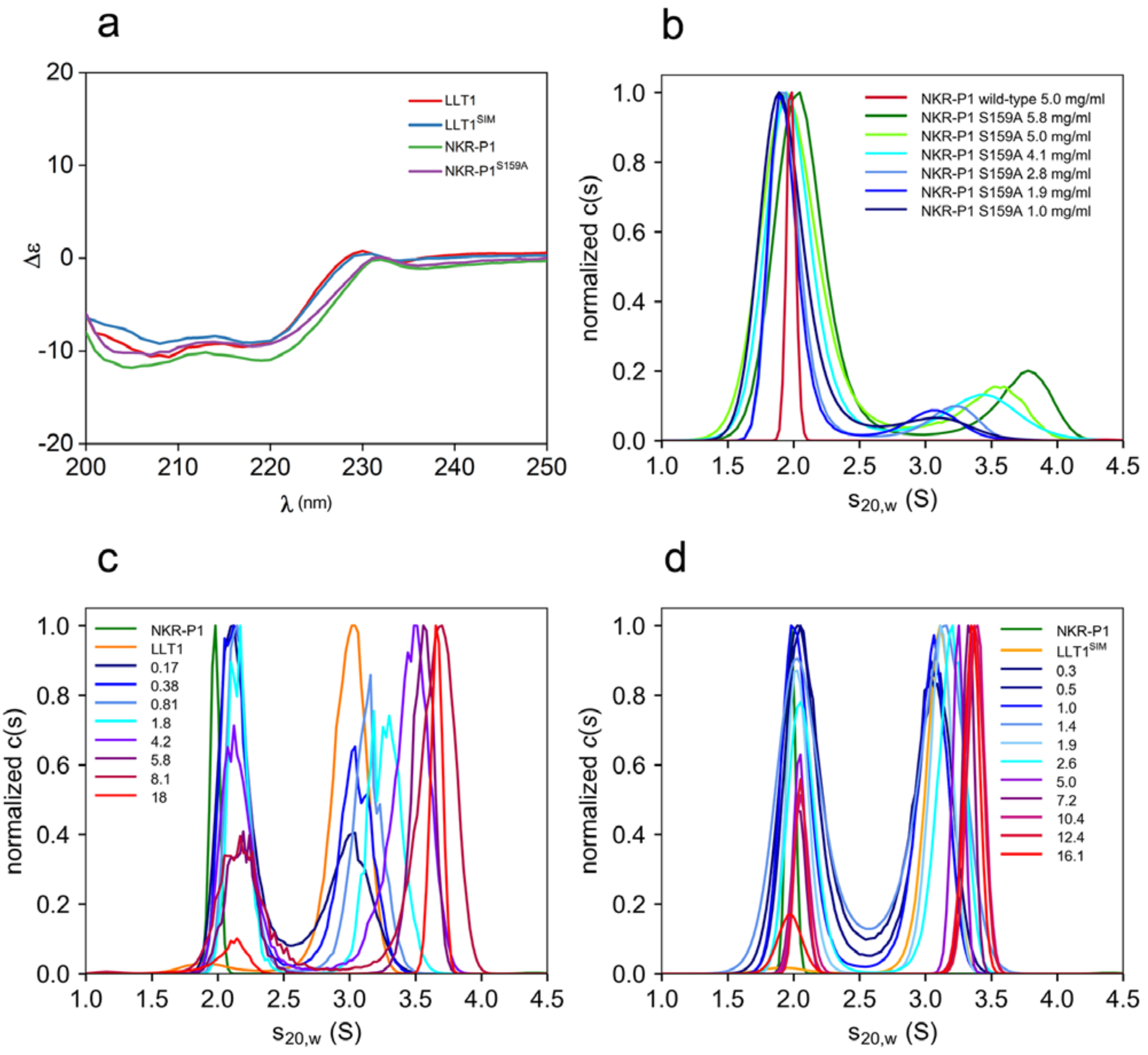

**Figure S4. NKR-P1 dimerization and NKR-P1:LLT1 complex formation observed in solution.** (**a**) CD spectra for wild-type LLT1 and NKR-P1 compared to the spectra of LLT1$^{SIM}$ and NKR-P1 S159A mutants used in sedimentation velocity analytical ultracentrifugation experiments. (**b**) Normalized continuous c(s) distributions of sedimentation coefficient transformed to standard conditions ($s_{20,w}$) for wild-type NKR-P1 (red) and increasing concentrations of NKR-P1 S159A mutant (dark blue to green). The shift of the distributions to higher S values corresponds to NKR-P1 oligomerization. (**c**) Normalized continuous c(s) distributions of sedimentation coefficient transformed to standard conditions ($s_{20,w}$) for free NKR-P1 (green, 5 mg/ml), free LLT1 (orange, 3.5 mg/ml) and their equimolar mixtures at increasing concentrations (blue to red, total concentration in mg/ml). (**d**) Normalized continuous c(s) distributions of sedimentation coefficient transformed to standard conditions ($s_{20,w}$) for free NKR-P1 (green, 5 mg/ml), free LLT1$^{SIM}$ (orange, 0.7 mg/ml) and their equimolar mixtures at increasing concentrations (blue to red, total concentration in mg/ml). The shift of the distributions to higher S values in (c) or (d) corresponds to NKR-P1:LLT1 or NKR-P1:LLT1$^{SIM}$ complex formation with fast kinetics, respectively.

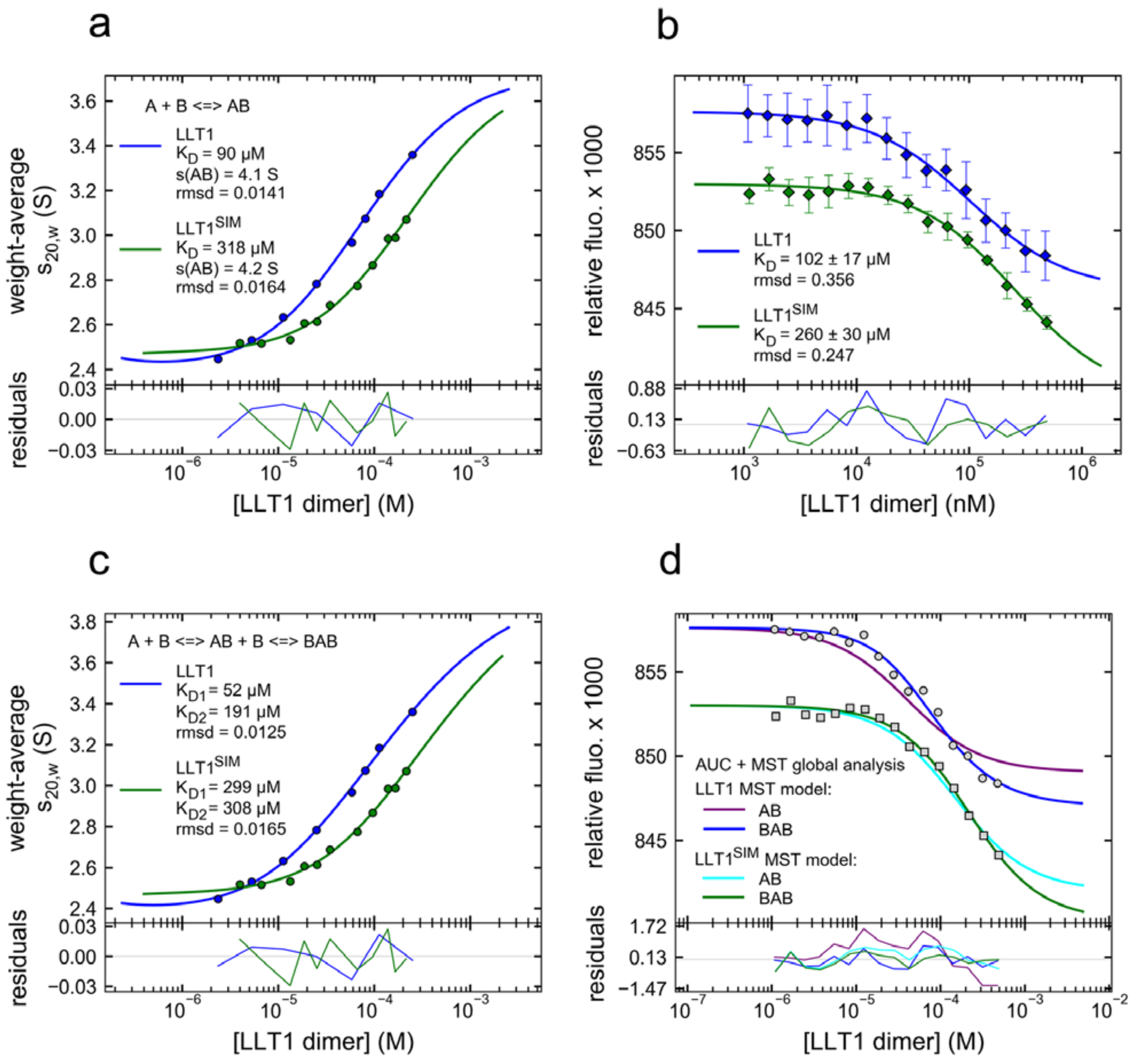

**Figure S5. NKR-P1:LLT1 interaction analyzed by analytical ultracentrifugation and microscale thermophoresis.** (**a**,**c**) Binding isotherms constructed by integrating the distributions shown in Fig. S4c,d over the whole range and plotting the resulting $s_{20,w}$ values against the LLT1 dimer concentration were best-fit with A+B⇔AB model where A is the LLT1 dimer and B is the NKR-P1 monomer in (a) or with A+B⇔AB+B⇔BAB model in (c). (**b**) Microscale thermophoresis data from three independent measurements were averaged and best-fit with A+B⇔AB model where A is the LLT1 dimer and B is the NKR-P1 monomer. (**d**) Global analysis of sedimentation velocity binding isotherms (c) and microscale thermophoresis data (b). Sedimentation parameters were fixed to the previously best-fit values, and the fluorescence isotherms were fit with either the AB or the BAB model. While for the LLT1$^{SIM}$ the AB model describes the data almost equally well as the BAB model, for wild-type LLT1, the AB model shows poor fit. The LLT1$^{SIM}$ data in (b) and (d) were purposely offset 0.004 relative fluorescence units for clarity.

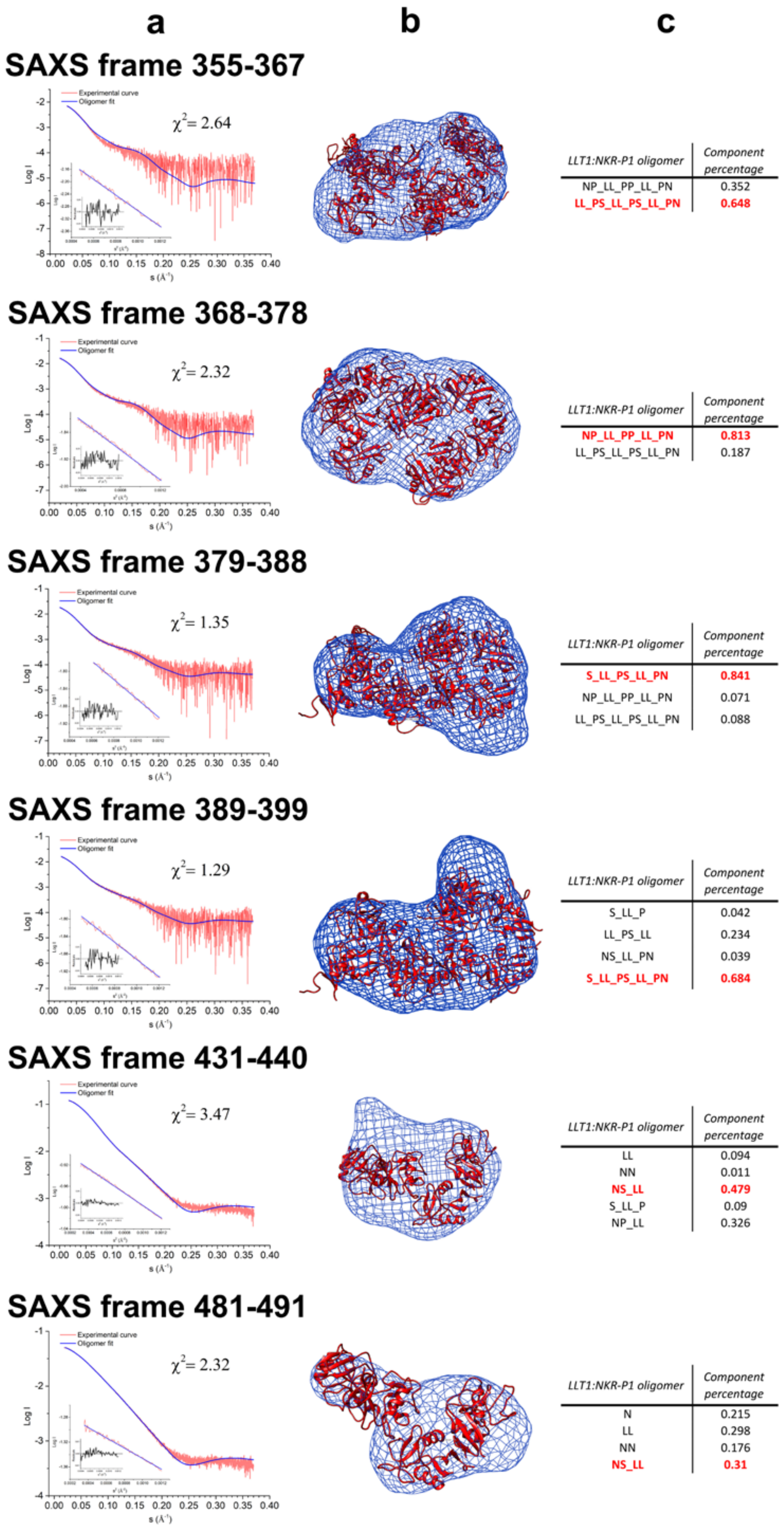

**Figure S6. OLIGOMER analysis of selected intervals of SEC-SAXS data observed for NKR-P1:LLT1 complexes.** (**a**) SAXS scattering curve in logarithmic scale (red) with the best-fit scattering curve calculated using OLIGOMER (blue). Guinier plot is shown in the lower-left corner, and the quality of the OLIGOMER fit is shown as a $\chi^2$ value. (**b**) Averaged and filtered DAMMIF envelopes (blue mesh) best-fit with the structure of the most abundant component of NKR-P1:LLT1 complex (red ribbon diagram) used to calculate the OLIGOMER scattering curve (highlighted in red also in (c)). (**c**) List of the superposed structures and their percentage abundance used to calculate the OLIGOMER scattering curve. **L** denotes the LLT1 monomer, **N** the NKR-P1 monomer, and **P** and **S** denote the NKR-P1 monomer engaged in the primary or secondary interaction mode with neighboring LLT1, respectively.

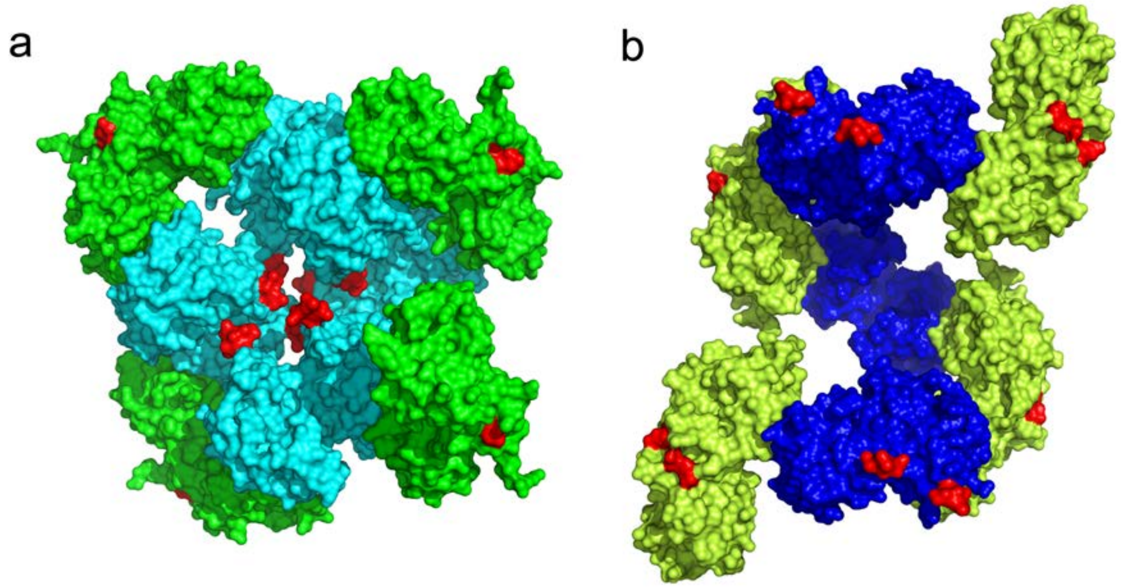

**Figure S7. Theoretic *in silico* models of the NKR-P1:LLT1 complex oligomeric assemblies, based on the propagation of only a single interaction mode, are biologically improbable.** (**a**) Theoretic oligomer based on the primary interaction mode only. LLT1 is in green, NKR-P1 in cyan. (**b**) Theoretic oligomer based on the secondary interaction mode only. LLT1 is in lemon, NKR-P1 in blue. The models were created by applying symmetry operations to parts of the NKR-P1:LLT1 complex crystal structure to artificially keep and propagate only the primary or the secondary binding mode. The first three N-terminal residues of the proteins are highlighted in red to highlight their orientation. In both (a) and (b), positions of the N-termini are incompatible with the physiological topology of the complex, i.e., NKR-P1 receptor on NK cell membrane and LLT1 ligand on the surface of another cell.

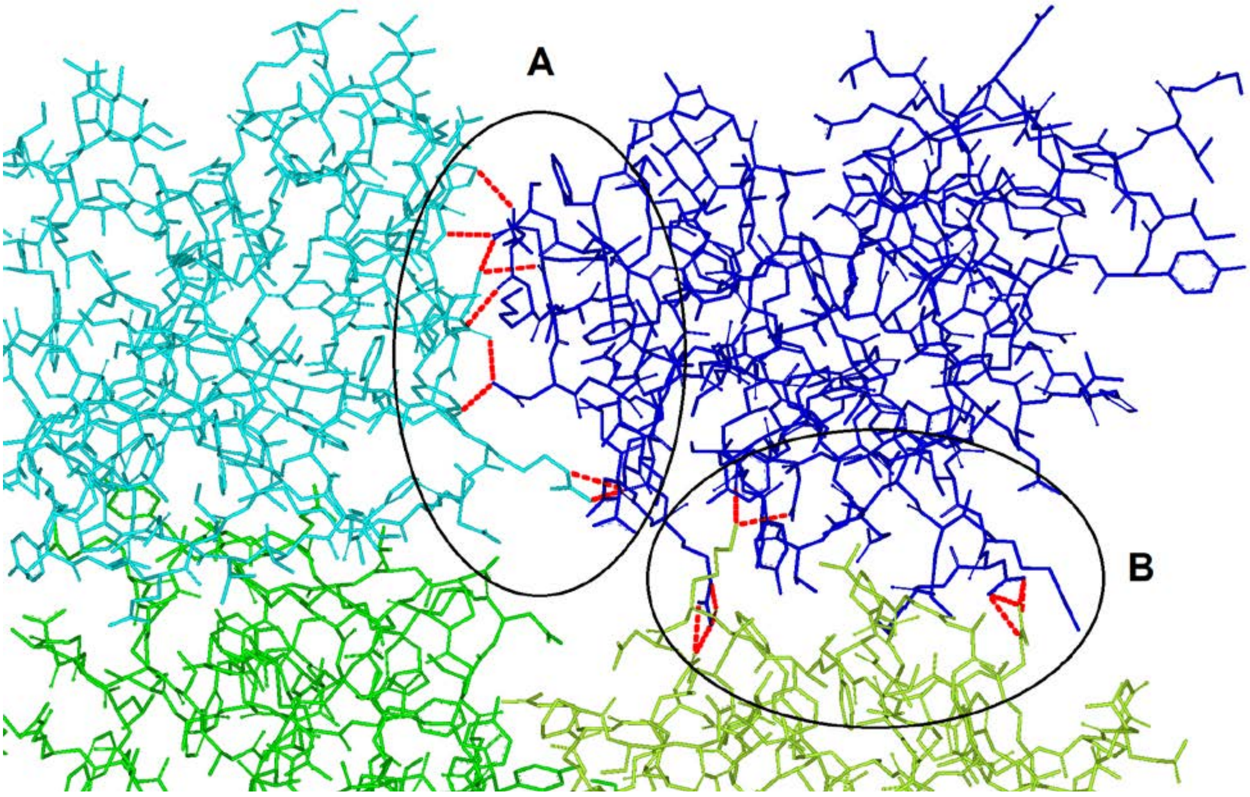

**Figure S8. NKR-P1 accessory interaction interface stabilizes the NKR-P1:LLT1 complex in primary and secondary binding mode.** A detail of the NKR-P1:LLT1 complex structure showing NKR-P1 and LLT1 interacting in the primary binding mode (cyan and green, respectively) and the secondary binding mode (blue and lemon, respectively). The accessory interaction interface (A) between the two neighboring NKR-P1 molecules ligated in the primary and secondary mode (cyan and blue, respectively) has a similar size and number of hydrogen bonds as the secondary interaction mode of the complex (B), thus stabilizing this complex's arrangement. This type of NKR-P1:NKR-P1 contact is not present in the theoretic models depicted in Figure S7a,b above.

**Table S1. Hydrogen bonds in the dimerization interface of NKR-P1, in the primary and secondary binding modes of the NKR-P1:LLT1 complex, and within the mutual contact of NKR-P1 bound in primary and secondary mode.** Water-mediated protein-protein hydrogen bonds are not shown.

| **NKR-P1 – chain A** | **NKR-P1 – chain B** | **Hydrogen bonds** | **Distance [Å]** |
|---|---|---|---|
| *helix α1-centered dimerization (NKR-P1_glyco: chain A, NKR-P1_glyco: chain B)* | | | |
| Ser127 | Ser123 | A/Ser127 O:B/Ser123 Oγ | 2.8 |
| Ile168 | Thr124 | A/Ile168 O:B/Thr124 Oγ1 | 3.1 |
| Glu213 | Glu126 | A/Glu213 N:B/Glu126 Oε2 | 2.8 |
| Ser123 | Ser127 | A/Ser123 Oγ:B/Ser127 O | 2.7 |
| Thr124 | Ile168 | A/Thr124 Oγ1:B/Ile168 O | 3.1 |
| Glu126 | Glu213 | A/Glu126 Oε2:B/Glu213 N | 2.8 |
| Asn157 | NAG501 | A/Asn157 Nδ2:B/NAG501 O7 | 3.0 |
| Ser159 | NAG501 | A/Ser159 Oγ:B/NAG501 O3 | 3.3 |
| NAG501 | Asn157 | A/NAG501 O7:B/Asn157 Nδ2 | 2.9 |
| NAG501 | Ser159 | A/NAG501 O4:B/Ser159 Oγ | 3.3 |
| NAG501 | Ser161 | A/NAG501 O4:W615/W632:B/Ser161 Oγ | 4.8 |
| **LLT1** | **NKR-P1** | **Hydrogen bonds** | **Distance [Å]** |
| *Primary mode (LLT1: chain B, NKR-P1: chain D)* | | | |
| Ser129 | Lys148 | Ser129 Oγ:Lys148 O | 2.6 |
| Asp130 | Ala149 | Asp130 N:Ala149 O | 3.0 |
| Glu162 | Tyr201 | Glu162 Oε1:Tyr201 OH | 2.5 |
| Arg175 | Asp183 | Arg175 Nη1:Asp183 Oδ2 | 2.9 |
| Arg175 | Asp183 | Arg175 Nη2:Asp183 Oδ2 | 3.2 |
| Arg175 | Glu200 | Arg175 N:Glu200 Oε2 | 3.0 |
| Tyr177 | Asp183 | Tyr177 OH:Asp183 N | 3.2 |
| Tyr177 | Ser199 | Tyr177 OH:Ser199 O | 2.6 |
| Glu179 | Arg181 | Glu179 Oε1:Arg181 Nη1 | 3.1 |
| Glu179 | Arg181 | Glu179 Oε1:Arg181 Nη2 | 3.0 |
| Glu179 | Tyr198 | Glu179 N:Tyr198 OH | 3.2 |
| *Secondary mode (LLT1: chain A, NKR-P1: chain C)* | | | |
| Asn120 | Arg181 | Asn120 Oδ1:Arg181 Nη1 | 3.0 |
| Asn120 | Arg181 | Asn120 Oδ1:Arg181 Nη2 | 3.0 |
| Arg153 | Asp147 | Arg153 Nε:Asp147 Oδ1 | 3.2 |
| Arg153 | Asp147 | Arg153 Nη2:Asp147 Oδ2 | 2.8 |
| Lys169 | Arg181 | Lys169 O:Arg181 Nη1 | 3.1 |
| Lys169 | Arg181 | Lys169 O:Arg181 Nε | 3.2 |
| Lys169 | Ser199 | Lys169 Nζ:Ser199 O | 2.6 |
| Lys169 | Glu200 | Lys169 Nζ:Glu200 Oε1 | 3.1 |
| **NKR-P1 – chain D** | **NKR-P1 – chain $C_{sym}$** | **Hydrogen bonds** | **Distance [Å]** |
| *Primary-secondary mode (NKR-P1: chain D, NKR-P1: chain $C_{sym}$)* | | | |
| Gln100 | Ser161 | Gln100 N:Ser161 O | 2.9 |
| Gln99 | Glu162 | Gln99 Nε2:Glu162 O | 3.0 |
| Asn143 | Asn174 | Asn143 Nδ2:Asn174 O | 3.3 |
| Asn143 | Asn174 | Asn143 Nδ2:Asn174 Oδ1 | 2.8 |
| Asn143 | Asn176 | Asn 143 O:Asn 176 Nδ2 | 3.4 |
| Ile145 | Asn176 | Ile 145 O:Asn 176 Nδ2 | 3.0 |
| Arg146 | Ile180 | Arg146 Nη1:Ile 180 O | 2.8 |

**Table S2. Interaction interface residues conserved among homologous NK cell CTL receptor:ligand complexes.** Pairs of interacting amino acid residues conserved in the primary interaction mode of human NKR-P1:LLT1, murine NKR-P1B:Clrb, and human NKp65:KACL complexes' crystal structures are listed, together with their mutual distances.

| **NKR-P1:LLT1** | **Distance [Å]** | **NKR-P1B:Clrb** | **Distance [Å]** | **NKp65:KACL** | **Distance [Å]** |
|---|---|---|---|---|---|
| D/Ser199 O: B/Tyr177 OH | 2.6 | U/Ser199 O: I/Tyr183 OH | 3.0 | *Phe160 in place of Tyr* | - |
| D/Glu205 O: B/Tyr165 OH | 3.6 | U/Asp205 O: I/Tyr171 OH | 2.6 | *Phe148 in place of Tyr* | - |
| D/Asp183 Oδ2: B/Arg175 Nη1 | 2.9 | U/Ser188 Oγ: I/Arg181 Nη1 | 3.3 | B/Ser171 Oγ: A/Arg158 Nη1 | 2.5 |
| D/Asp183 Oδ2: B/Arg175 Nη2 | 3.2 | U/Ser188 Oγ: I/Arg181 Nη2 | 3.1 | B/Ser171 Oγ: A/Arg158 Nη2 | 3.1 |
| - | - | U/Ser199 Oγ: I/Arg181 Nη1 | 3.6 | B/Ser182 Oγ: A/Arg158 Nη2 | 3.4 |
| *D/Ser199 O: to Lys169 Nζ in secondary mode* | *2.6* | U/Ser199 O: I/Arg181 Nη1 | 3.4 | B/Ser182 O: A/Arg158 Nη2 | 3.1 |

**Table S3. Comparison of the previously proposed NKR-P1:LLT1 binding model with the crystal structure.** The table lists point mutations of LLT1 and NKR-P1 residues that had either a detrimental or moderate negative effect on the binding of its partner as determined by SPR analyses in the previous works of Kamishikiryo *et al.* and Kita *et al.*[2, 3]. The amino acid interaction pairs proposed in these studies are listed at the bottom. The presence or absence of such residue or interaction pair within the primary or secondary binding mode interface in the herein reported NKR-P1:LLT1 complex crystal structure is indicated with upper indices (P – primary, S – secondary, - – absence).

| LLT1 | NKR-P1 |
|---|---|
| ***Detrimental effect on binding*** | |
| Lys169Glu $^{P/S}$ | Glu162Arg $^{-/-}$ |
| Arg175Glu $^{P/-}$ | Asp183Arg $^{P/S}$ |
| Arg180Glu $^{P/-}$ | Tyr198Ala $^{P/S}$ |
| Lys181Glu $^{P/-}$ | Tyr201Ala $^{P/S}$ |
| | Glu205Arg $^{P/-}$ |
| ***Moderate effect on binding*** | |
| Tyr165Ala $^{P/S}$ | Arg181Glu $^{P/S}$ |
| Asn167Ala $^{P/-}$ | Glu186Arg $^{-/-}$ |
| ***Proposed LLT1:NKR-P1 interaction pairs*** | |
| Lys169:Glu205 $^{-/-}$ | |
| Arg175:Glu200 $^{P/-}$ | |
| Glu179:Ser193/Thr195 $^{P/-}$ | |
| Tyr177:Tyr198 $^{P/-}$ | |
| Tyr165:Phe152 $^{-/-}$ | |
